# Age-dependent normalisation functions for T-lymphocytes in healthy individuals

**DOI:** 10.1101/2021.12.01.470754

**Authors:** Juliane Schröter, José A. M. Borghans, W. Marieke Bitter, Jacques J. M. van Dongen, Rob J. de Boer

**Affiliations:** Theoretical Biology & Bioinformatics, Utrecht University, Utrecht, Netherlands; Center for Translational Immunology, University Medical Center Utrecht, Utrecht, Netherlands; Department of Immunology, Leiden University Medical Center, Leiden, Netherlands

## Abstract

Lymphocyte numbers naturally change through age. Normalisation functions to account for this are sparse, and mostly disregard measurements from children in which these changes are most prominent. In this study, we analyse cross-sectional numbers of mainly T-lymphocytes (CD3^+^, CD3^+^CD4^+^ and CD3^+^CD8^+^) and their subpopulations (naive and memory) from 673 healthy Dutch individuals ranging from infancy to adulthood (0-62 years). We fitted the data by a delayed exponential function and received parameter estimates for each lymphocyte subset. Our modelling approach follows general laboratory measurement procedures in which absolute cell counts of T-lymphocyte subsets are calculated from observed percentages within a reference population that is truly counted (typically the total lymphocyte count). Consequently, we receive one set of parameter estimates per T-cell subset representing both the trajectories of their counts and percentages. We allow for an initial time delay of half a year before the total lymphocyte counts per *µ*l of blood start to change exponentially, and we find that T-lymphocyte trajectories tend to increase during the first half a year of life. Thus, our study provides functions describing the general trajectories of T-lymphocyte counts and percentages of the Dutch population. These functions provide important references to study T-lymphocyte dynamics in disease, and allow one to quantify losses and gains in longitudinal data, such as the CD4^+^ T-cell decline in HIV-infected children, and/or the rate of T-cell recovery after the onset of treatment.

## Introduction

Lymphocyte numbers experience dynamical changes over a lifetime – known as a hallmark of a maturing and subsequently ageing immune system [Wade and Ades, 1994, Hulstaert et al., 1994, Bains et al., 2009, Valiathan et al., 2016, van den Heuvel et al., 2017]. These dynamics are more prominent in childhood than in adulthood, as a child’s immune system is maturing and has not yet approached steady state. These natural changes over time have to be considered while determining the immunological health status of an individual.

To define an individual’s immunological health status, cell counts and percentages of various lymphocyte subsets are measured and compared to reference values of healthy individuals [Picat et al., 2013, Ásbjörnsdóttir et al., 2016, Lewis et al., 2017, Salzmann-Manrique et al., 2018, Schröter et al., 2020]. For this reason, lymphocyte data of healthy individuals have been presented in contiguous age-categories, combining lymphocyte measurements from several months to several years into bins [Shearer et al., 2003, Comans-Bitter et al., 1997, van Gent et al., 2009, Lawrie et al., 2015, Blanco et al., 2018, Van Dongen et al., 2019]. These age-intervals allow clinicians to classify the health status of an individual at a given age.

As these bins often cover a broad age range and the lymphocyte numbers themselves are variable, continuous and gradual changes in lymphocytes over time during a disease – such as the decline in CD4+ T cell numbers during HIV infection, and/or their recovery after the onset of treatment – are difficult to quantify. Instead of binning the data into age categories, one can also define continuous normalisation functions as healthy references. By providing a finer time scale, such functions allow a better quantification of lymphocyte dynamics.

To our knowledge, only few publications present estimates describing lymphocyte data of healthy individuals in a continuous manner [Wade and Ades, 1994, Huenecke et al., 2008, Payne et al., 2020]. The often cited paper from Huenecke et al. [2008] describes the trajectories of several T-lymphocyte subsets and their subpopulations based on 100 healthy German individuals by fitting an exponential function, decaying towards an asymptote. Unfortunately, this dataset only includes 14 children under the age of one year, which is the time frame during which one expects major dynamical changes. Moreover, this paper neglects an initial increase within the first half a year of age as observed by longitudinal analyses and in large cohort studies [de Vries et al., 1998, van den Heuvel et al., 2017, Li et al., 2020]. Additionally, Huenecke et al. [2018] provides neither parameters for the total lymphocyte counts, nor for the CD4^+^ and CD8^+^ percentages. Percentages usually show less variability than the cell counts per *µ*l of blood [Goicoechea and Haubrich, 2005], and they are easier and cheaper to obtain through flow cytometry analysis [Comans-Bitter et al., 1997]. Thus, percentages are more commonly reported than counts. As percentages cannot replace the information retrieved from count data, normalisation functions for both counts and percentages are required.

To extent on this and to obtain a much more detailed description of the T-lymphocyte dynamics during the first year of life, we fitted cross-sectional age-matched total lymphocyte and T-lymphocyte measurements from 673 healthy Dutch individuals (ranging from 0 to 62 years), including 268 individuals under the age of one year. As a statistical improvement, our methods follow laboratory measurement procedures, where most cell counts are calculated from observed percentages within a truly counted reference population. We therefore fit percentage data, while receiving parameters describing the cell count data. Thus, we provide new standard functions expressing both absolute and relative data for several T-lymphocyte subsets and their subpopulations of the Dutch population.

## Method

### Data

We accessed the age-matched raw data of two published Dutch cohorts: Dutch I (presented by bullets (•) in all figures) [Comans-Bitter et al., 1997] and Dutch II (presented by triangles (▲) in all figures) [van Gent et al., 2009]. Both publications presented their lymphocyte data in age-categories. The data presented in Dutch I are typically used as standard reference intervals for the Netherlands and Europe [Shearer et al., 2003, Huenecke et al., 2008, Payne et al., 2020]. For Dutch I, we had measurements from 426 individuals including total lymphocyte counts (TLC) as well as the counts and percentages of the CD3^+^, CD3^+^CD4^+^ and CD3^+^CD8^+^ subsets within the total lymphocyte gate. For Dutch II, we had measurements from 112 children and adolescents including total lymphocyte counts as well as the counts and percentages of the CD3^+^CD4^+^, CD3^+^CD8^+^, naive CD4+ (CD27^+^ CD45RO^*−*^ CD4^+^), memory CD4+ (CD45RO^+^ CD4^+^), effector CD4+ (CD27^*−*^ CD45RO^*−*^ CD4^+^), naive CD8+ (CD27^+^ CD45RO^*−*^ CD8^+^), memory CD8+ (CD45RO^+^ CD8^+^), and effector CD8+ (CD27^*−*^ CD45RO^*−*^ CD8^+^) subsets. To cover the naive, memory, and effector subpopulations until late adulthood, we accessed a third dataset (Dutch III; presented by squares (▪) in all figures) with an additional 135 individuals [Veel et al., 2018]. For Dutch III, percentages and count data are given for the CD4+ and CD8+ T-cell naive, memory, and effector subpopulations, but only count data is provided for CD4+ and CD8+ T-cell subsets. Even though the cohorts were collected a decade apart in different laboratories, their quantifications of count data were based on similar methods (in Dutch I with a Coulter Counter model Z1, and in Dutch II/III with a Cell-Dyn Sapphire™ Hematology Analyzer) and we do not observe major differences by visual inspection of cross-sectional illustrations of the data (Figure 1-3). Thus, we combined the three cohorts for our analysis and refer to them from now on as the Dutch cohorts (see also Table S1).

**Figure 1:**
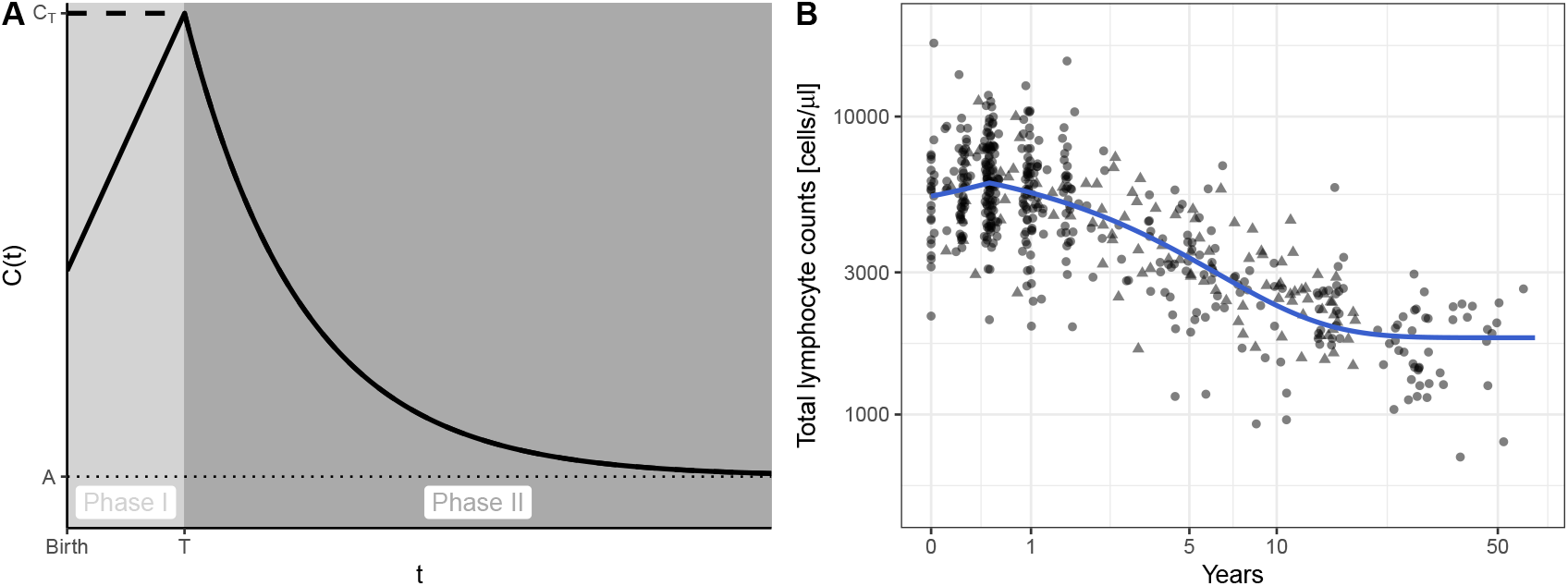
Lymphocyte trajectory follows a delayed exponential decay towards and asymptote. (A) Schematic illustration of the mathematical model (Equation 1) with a flat (dashed line) and an increasing (solid line) initial linear phases (I) lasting until time *T*, followed by an exponential declining phase (II) from the value at *T* (C_T_) towards an asymptote *A* (dotted line). (B) Measured total lymphocyte cell counts/*µ*l blood over age (•: Dutch I, ▴: Dutch II) are shown along with the best fit of model (1) to the data (blue line).

### Model

#### Absolute cell counts

The natural decline of lymphocyte counts over age has previously been described phenomenologically by an exponential decay towards an asymptote [Huenecke et al., 2008]. To allow for a different initial phase, which has been observed in longitudinal data and large cohort studies [de Vries et al., 1998, van den Heuvel et al., 2017, Li et al., 2020], we extend the original model by defining two phases for the absolute cell counts per *µ*l blood, *C*(*t*), depending on time *t* in years (Figure 1A). The initial phase (Phase I) is described by a linear function which lasts until time *T*, and is succeeded by a second phase (Phase II) describing the exponential towards an asymptote as reported by Huenecke et al. [2008]:

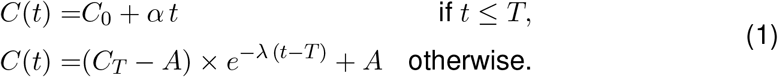

For any lymphocyte subset, *C*_0_ is the initial cell count per *µ*l blood at birth, which changes linearly until time *T* to a value *C*_*T*_ with a slope *α* (cells per year). From time *T* onwards the counts change exponentially from *C*_*T*_ at a rate *λ* (per year), and eventually approach an asymptote *A* (cells per *µ*l blood), which corresponds to the normal homeostatic level as seen in healthy adults. Depending on the initial value, *C*_0_, and the ultimate asymptote, *A*, our model (1) can describe either overall increasing or declining dynamics of different lymphocyte subsets. The exponential, *λ*, can be expressed as a half-life or doubling time, 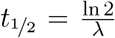, giving an indication for the time required to approach the homeostatic level, *A*, from the level at time *T, C*_*T*_. Since this is independent of the difference, *C*_*T*_ *− A*, in cell numbers, we also report the quantity *λ*(*C*_*T*_ *− A*), which is the initial slope after the early initial phase in terms of ‘cells per *µ*l per year’. Model (1) can be fitted to the absolute cell count data of any lymphocyte subset.

#### From percentages to absolute cell counts

In immunology, one is typically interested in both the absolute cell counts and the relative fractions (typically presented as percentages) of lymphocytes subsets. Since percentages are easier and cheaper to measure by flow cytometry than counting cells directly, the data representing the percentages are more commonly gathered than the cell counts of a given subset. In experiments, one typically counts some reference population in a *µ*l blood, like total lymphocytes, and estimates the fractions of various subsets within that reference population by flow cytometry. The cell counts of the subsets are then calculated by multiplying these observed fractions with the observed cell count of the reference population. We decided to fit the data according to this experimental procedure, and refrain from the conventional way of fitting model (1) directly to the, from now on referred, *“calculated”* cell counts.

Instead, we first fit model (1) to the cell counts of the reference population, *C*_*R*_(*t*) (in our case total lymphocytes). Keeping the estimated parameters of the reference population, we then fit the observed percentages, *P* (*t*), of a particular subset, by reversing the calculation of the cell counts from the measurement procedure, i.e.,

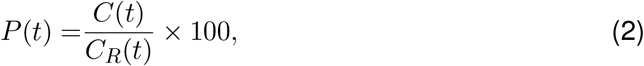

while fitting the parameters of model (1) to describe the unobserved cell counts, *C*(*t*), of the subset of interest (e.g., CD4+ T cells; see also Box 1). The advantages of this procedure are that (i) we stay close to the experimental measurement procedures, that (ii) we use the same parameters to describe the percentages and the cell counts of a particular subset, and that (iii) we can describe complicated trajectories of percentages that are decreasing and subsequently increasing over time with a very simple model. As a sanity check, we will show that fitting the “calculated” cell count data directly with model (1) provides similar estimates (see Figure S2, Table S3).

#### Algorithm and confidence intervals

To fit the models to the data, we use the Grind wrapper that is based on the `FME` package in R (version 4.0.2) [R Development Core Team, 2003, Soetaert and Petzoldt, 2010]. Parameter estimates are obtained based on the Levenberg-Marquardt algorithm by minimising the sum of squared residuals. We fit the logarithmically transformed data to account for the measurement variance of several magnitudes over age (particularly in the count data) and to receive normally distributed residuals. Parameter estimates are constrained to be positive, and are tested on being significantly different from zero by the F-test at a significance level of 0.05 (as provided by the algorithm in the `FME` package). We additionally apply a bootstrapping method to obtain 95% confidence intervals for each parameter estimate, by resampling the data 1000 times. The 95% confidence ranges for the parameters are displayed in Table 1 and Table S3.

**Table 1:** Parameter estimates and their 95% confidence intervals to obtain fits for count data via model (1). The 95% confidence intervals for each parameter are received from bootstrapping the data 1000 times. The delay *T* was fixed to 0.5 years. The dimensions of the parameters are as follows, *C*_0_, *A*: cells per *µ*l, *α*: cells per *µ*l per year, *λ*: per year, (*C*_*T*_ *−A*)*λ*: cells per *µ*l per year, and *t* _½_: years.

| Subset | | $C_0$ | $\alpha$ | $\lambda$ | $A$ | $(C_T - A)\lambda$ | $t_{1/2}$ |
| --- | --- | --- | --- | --- | --- | --- | --- |
| Total lymphocyte count | Estimate | 5405.5 | 1165.1 | 0.221 | 1810.1 | 923.33 | 3.14 |
|  | 95% CI | [4794.0 - 5954.5] | [0.0 - 2785.9] | [0.174 - 0.287] | [1625.4 - 1986.5] | - | - |
| CD3+ T-cell count | Estimate | 3439.9 | 796.1 | 0.238 | 1287.0 | 607.13 | 2.91 |
|  | 95% CI | [3221.1 - 3654.2] | [299.2 - 1326.3] | [0.218 - 0.262] | [1242.4 - 1330.0] | - | - |
| CD4+ T-cell count | Estimate | 2424.7 | 490.1 | 0.380 | 788.9 | 714.72 | 1.82 |
|  | 95% CI | [2202.8 - 2622.8] | [0.0 - 987.6] | [0.341 - 0.427] | [755.5 - 821.5] | - | - |
| CD8+ T-cell count | Estimate | 963.7 | 211.8 | 0.156 | 386.5 | 106.56 | 4.44 |
|  | 95% CI | [866.0 - 1057.6] | [0.0 - 440.5] | [0.133 - 0.179] | [350.5 - 418.6] | - | - |
| naive CD4+ T-cell count | Estimate | 1733.1 | 0.0 | 0.162 | 428.1 | 211.41 | 4.28 |
|  | 95% CI | [1447.6 - 2122.1] | [0.0 - 0.0] | [0.104 - 0.299] | [340.5 - 548.4] | - | - |
| memory CD4+ T-cell count | Estimate | 140.4 | 167.2 | 0.600 | 332.1 | -64.86 | 1.16 |
|  | 95% CI | [49.2 - 228.3] | [0.0 - 454.4] | [0.004 - 10.476] | [309.9 - 999.9]* | - | - |
| effector CD4+ T-cell count | Estimate | 6.8 | 0.0 | 0.000 | 6.8 | 0.00 | - |
|  | 95% CI | [5.8 - 8.0] | - | - | [5.8 - 8.0] | - | - |
| naive CD8+ T-cell count | Estimate | 607.5 | 208.9 | 0.079 | 169.6 | 42.85 | 8.77 |
|  | 95% CI | [311.3 - 762.3] | [0.0 - 877.5] | [0.049 - 0.114] | [92.0 - 223.3] | - | - |
| memory CD8+ T-cell count | Estimate | 22.2 | 57.3 | 0.468 | 122.4 | -33.49 | 1.48 |
|  | 95% CI | [0.0 - 55.3] | [0.0 - 156.9] | [0.037 - 1.073] | [109.8 - 175.2] | - | - |
| effector CD8+ T-cell count | Estimate | 8.4 | 0.0 | 0.488 | 35.3 | -13.13 | 1.42 |
|  | 95% CI | [4.8 - 13.7] | [0.0 - 0.0] | [0.164 - 1.010] | [29.2 - 45.2] | - | - |

\* Parameter was constraint to an upper limit of 1000 cells/ $\mu\text{l}$

## Results

### No direct decline in lymphocyte counts after birth

We modelled lymphocyte dynamics from birth to 60 years of age with a simple delayed exponential function (Equation 1, Figure 1A). We accessed published reference lymphocyte data of two Dutch cohorts. Together they included measurements from 538 healthy individuals covering the full age range. Measurements are enriched for individuals under the age of one year (n=253), which allowed us to focus on T-lymphocyte dynamics in the very early phase of life, when changes are largest. Longitudinal data and large cohort studies [de Vries et al., 1998, van den Heuvel et al., 2017, Li et al., 2020] report an initial increase in lymphocyte counts during the first half a year of age (rather than the typically assumed exponential decline from birth onwards [Huenecke et al., 2008]). Visual inspection of our data (see Figure 1B, Figure 2, Figure 3) and linear regression analyses, resulting in positive slope estimates for the data covering the first year of life (see Table S2), support these observations. Consequently, we have extended the conventional exponential model [Huenecke et al., 2008] with an initial phase allowing for an early increase (Equation 1, Figure 1A).

**Figure 2:**
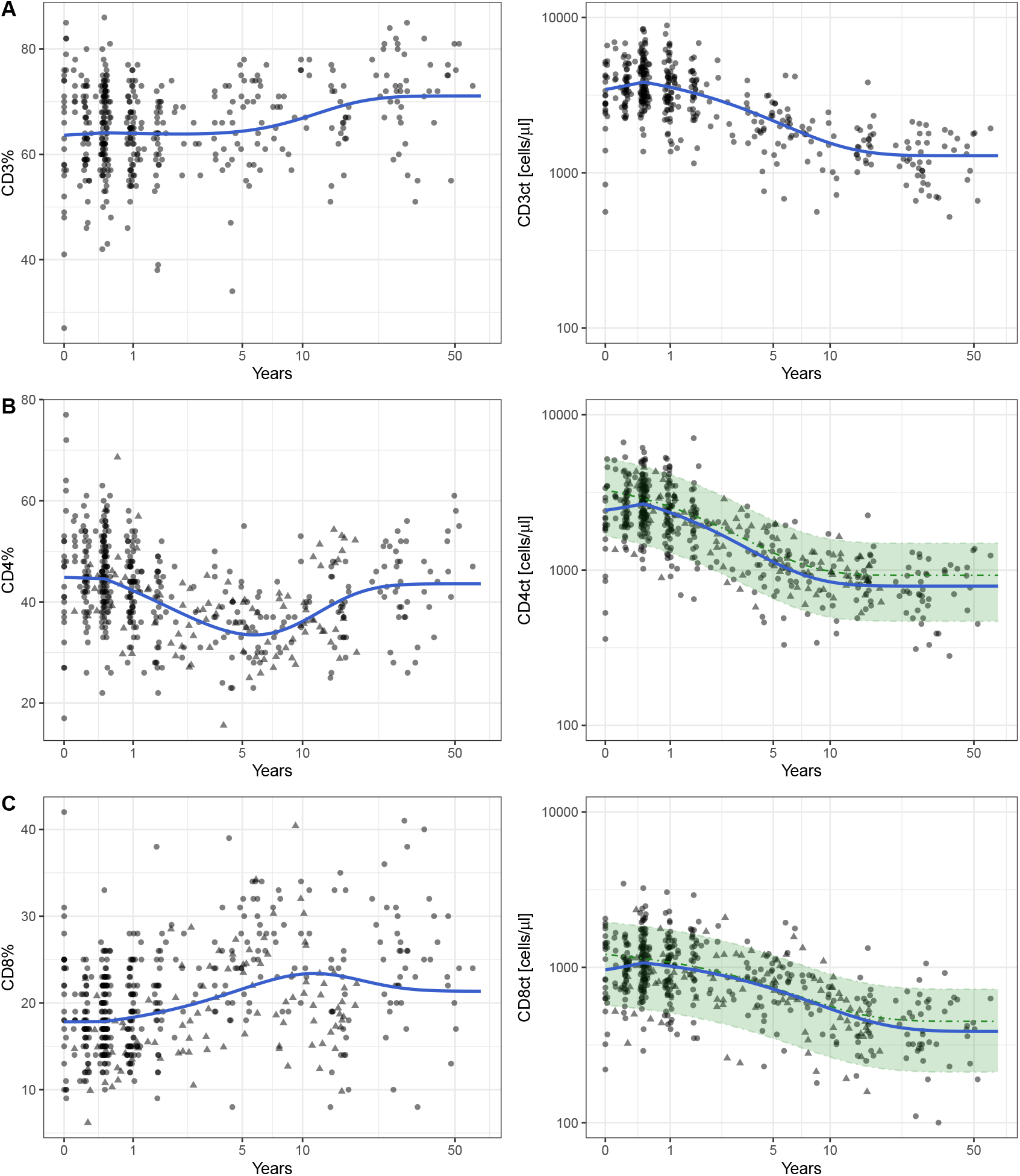
Trajectories of the T-lymphocyte subsets CD3+ (A), CD4+ (B), and CD8+ (C). The left panels illustrate the measured percentages and the right panels the calculated absolute counts. Data from Dutch I are represented by bullets (•), and data from Dutch II by triangles (▴). In the left panels, the blue solid lines represent the best fit of model (2). In the right panels, the blue solid lines represent the underlying trajectories for the calculated absolute count according to model (1). The parameter estimates listed in Table 1 are used. If available, the estimates from Huenecke et al. (2008) are presented in green with their 95% confidence range.

**Figure 3:**
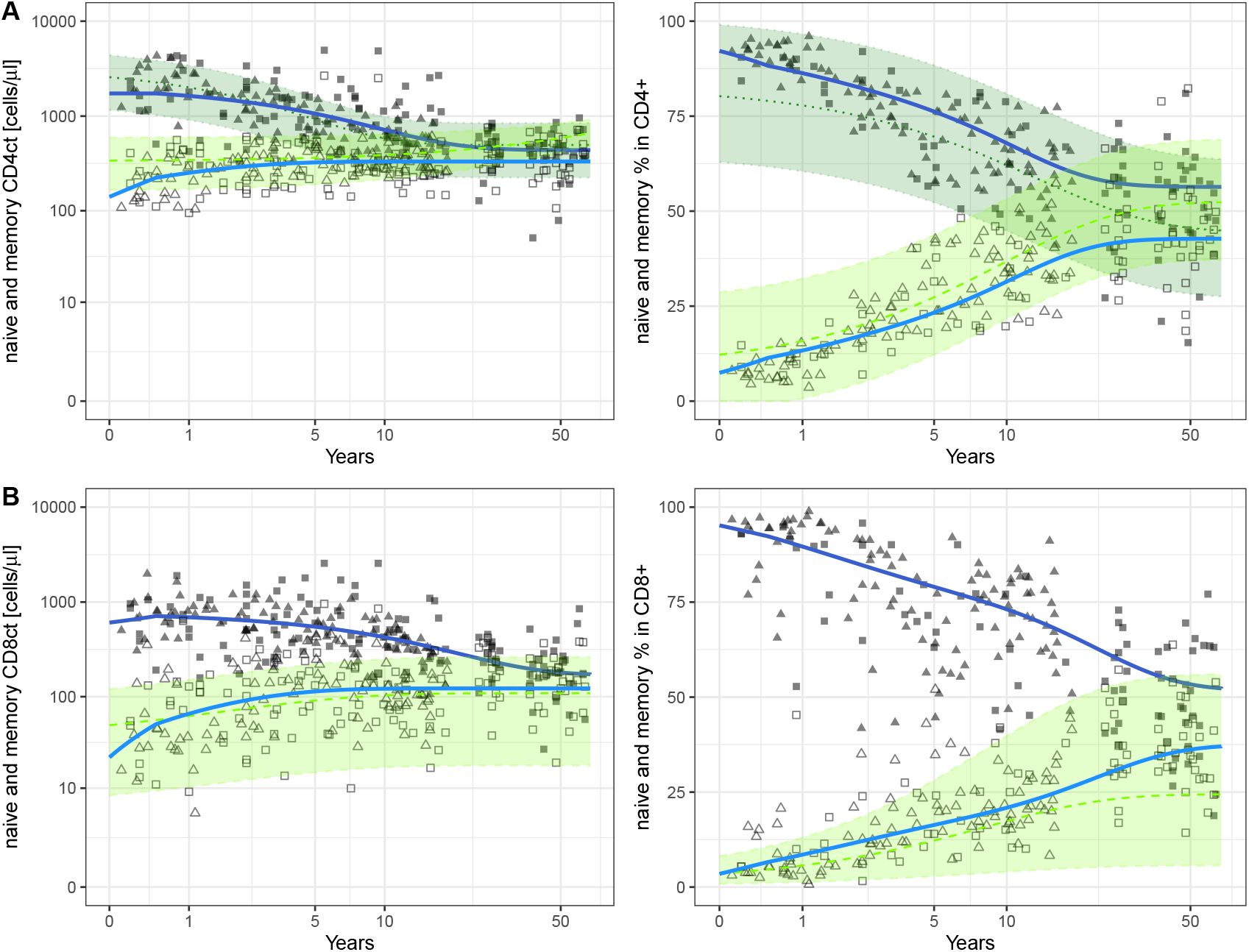
Trajectories of the naive and memory subpopulations of CD4+ (A) and CD8+ cells (B). The left panels give the absolute counts and the right panels the percentages. Data from Dutch II are represented by triangles (▴), and data from Dutch III by squares (▪). The naive subpopulation is represented by filled symbols, the memory subpopulations by open symbols. In the left panels, the blue solid lines represent the best fit of model (1) and the parameters reported in Table (1). In the right panels, the blue solid lines represent the underlying trajectories for the percentages according to model (2) where the sum of naive, memory and effector cell counts add up to 100%. The dark blue colour refers to the naive subpopulation the lighter blue to the memory subpopulations. The green dotted (naive) and dashed (memory) lines represent the estimates from Huenecke et al. (2008) and their 95% confidence range (green area). We noticed that Huenecke et al. [2008] described the percentages of the naive CD8+ subpopulation by an increasing exponential function, while our data (and fit) follows a decline. We excluded this prediction from Huenecke et al. [2008] in panel (B) (i.e., the missing green range for naive CD8+ counts), because we assume this to be a mistake (since the percentage of naive T cells should decline to general immunological knowledge).

### Fitting total lymphocyte counts (TLC) as a reference population

An attempt to freely fit all five parameters of the model (Equation 1) to the cross-sectional total lymphocyte counts (TLC) resulted in high correlations of the parameters describing the first phase (*C*_0_, *α* and *T*). Thus, these 3 parameters are no identifiable from our data. We therefore decided to fix *T* to the in literature reported minimum value of *T* =0.5 years [de Vries et al., 1998, van den Heuvel et al., 2017, Li et al., 2020], which reduces the number of free parameters in the model (Equation 1) from five to four. The best fit of the TLC data results in an linearly increasing phase over the first half a year from *C*_0_=5406 cells/*µ*l to *C*_*T*_ =5988 cells/*µ*l, before the TLC starts to decline exponentially at a rate of *λ*=0.221 per year towards an asymptotic level of *A*=1810 cells/*µ*l (Figure 1B, Table 1). Identifying the slope *α* of the initial phase remains difficult (see Table 1), because *α* correlates with the initial value *C*_0_, and *α* is not significantly different from zero. This is due to the wide spread of the cross-sectional data suggesting inter-individual variabilities. Given the good evidence for an initial increase in [de Vries et al., 1998, van den Heuvel et al., 2017, Li et al., 2020], we continued with the non-zero *α*= 1165 cells/*µ*l, as we aimed to provide a TLC reference function for the Dutch population including the additional knowledge of an early increase. The parameters of the best fit implemented into model (1) result in a smooth trajectory describing the overall dynamics of the TLC data reasonably well (Figure 1B, Table 1). This smooth TLC trajectory served as the reference to obtain the following estimations for the trajectories of the CD3+, CD4+ and CD8+ T-cell subsets via model (2).

### Box 1

**Example calculations**

Model (1) provides normalisation functions for count data with the parameters pre-sented in Table 1. For example, for the reference population TLC and CD4 count (CD4ct) in cells per *µ*l:

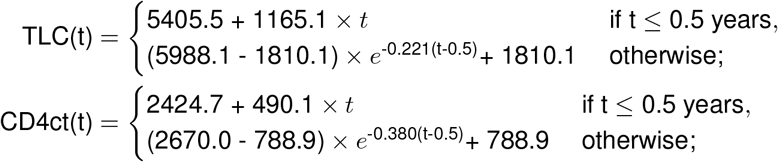

Model (2) provides normalisation functions for the percentages. For example, for CD4%:

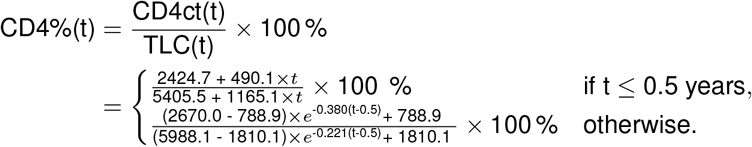

We first fit the function TLC(t) to the TLC data, and subsequently fit the function CD4%(t) to the percentage data by estimating the four free parameters of the CD4ct(t) function.

### Fitting non-monotonic percentages to receive cell count predictions

In contrast to the TLC, absolute counts of lymphocyte subsets, such as CD3+, CD4+, CD8+ T cells are typically calculated from the observed fraction within the lymphocyte gate, i.e., they are not directly counted. While the calculated count data obey the trajectories described by model (1), the observed percentages of several lymphocyte subsets change in a more complicated manner over time, with an increasing and a decreasing phase (Figure 2 left panels), which is more challenging to model. By following typical experimental procedures, and using model (2) with the TLC as a reference, we can describe these non-monotonic trajectories with a rather simple function (Figure 2 left panels). Thus, each trajectory can be estimated by four additional free parameters (*C*_0_, *α, λ*, and *A*), predicting the corresponding T-cell counts by inserting them into model (1) (see Box 1).

Taking the CD4% as an example (Figure 2B), one can recognise the an almost flat phase over the first half a year at a value of 45%, before the CD4% drops by about 10 percent points to reach a nadir around the age of five years. Afterwards, the CD4% increases again to approach a plateau – at a level similar to the one observed at birth – in adulthood. Thus, a “wavy” CD4% trajectory is described, which confirms a previous observation [Valiathan et al., 2016]. The CD3% and CD8% both expand from birth onwards, and also approach a homeostatic level around the age of 30 (Figure 2A/C). Hence, by staying close to experimental procedures we model trajectories of the percentages that have more complicated dynamics than was previously appreciated.

Although our predictions for the calculated count data are based upon fitting the observed percentages, they are in good agreement with the calculated CD3+, CD4+, and CD8+ T-cell counts (Figure 2 right panels). Additionally, the trajectories predicted by our procedure of estimating observed data only, are in excellent agreement with the trajectories that are obtained by estimating the calculated count directly with model (1) (Figure S1). Thus, the normalisation functions obtained with either method (Table 1 or Table S3) are very similar.

### Fitting complementary T-cell subpopulations

Lastly, we provide estimates for the naive, effector and memory subpopulations of CD4+ and CD8+ T cells (Figure 3, Table 1). Those subpopulations are usually reported as fractions of either CD4+ or CD8+ T cells and sum up to almost 100%. Unfortunately, these reference populations are typically not truly counted, which precluded us from using model (2). We therefore fitted the calculated count data using model (1) (Figure 3 and Figure S2 left panels, Table 1). The three trajectories (naive, memory and effector subpopulations) are then considered to sum up to 100%, from which we calculated trajectories for the percentages of the subpopulations according to model (2) (Figure 3 and Figure S2 right panels).

Our results (in blue) fit the general trajectories of the data reasonably well (in grey). At birth the CD4+ and CD8+ T-cell pools are largely composed of naive T cells (80-90%; Figure 3). Naive T-cell counts decline about 4-fold over 50 years, i.e., from 1733 to 438 cells per *µ*l for CD4+ and from 608 to 170 cells per *µ*l for CD8+ T cells (Figure 3 and Table 1). The memory compartments show opposite dynamics and increase over time towards an asymptote (Figure 3). Overall our trajectories are in good agreement with those reported by Huenecke et al. [2008] (in green). Note that Huenecke et al. [2008] describe the memory CD4+ and naive CD8+ T-cell counts by a linear model, whereas we also consider the effect of homeostasis in the subpopulations.

## Discussion

We present a new mathematical approach to describe the natural dynamics in lymphocyte numbers continuously. By providing parameters for various T-lymphocyte subsets (i.e., CD3+, CD4+, CD8+ T cells), and their subpopulations (naive and memory), reference functions for both their absolute counts and percentages can be formulated. Based on an extensive dataset, including age-matched cross-sectional lymphocyte measurements of 673 healthy Dutch individuals (covering particularly the dynamic first year of life), we are able to describe all trajectories phenomenologically with a bi-phasic model. We allowed for an initial phase of half a year in which lymphocyte subsets can increase, followed by a second phase which describes an exponential that approaches an asymptote corresponding to the homeostatic adult level. Except for memory T cells, this homeostatic level is below the values at birth. Our approach of fitting percentages to the ratio of two simple exponential models allowed us to simultaneously describe the counts and percentages of all populations of interest, delivering non-monotonic trajectories of the CD4+ and CD8+ T-cell percentages. We provide a consistent set of parameters describing both percentages and their dependent counts.

We extended the model of a similar previous publication [Huenecke et al., 2008] with an initial phase to relax the previous assumption that the TLC declines from birth onwards. Longitudinal data, as well as large cross-sectional studies, demonstrated that cell numbers per *µ*l blood increase during the first half a year of life [de Vries et al., 1998, van den Heuvel et al., 2017, Li et al., 2020]. We checked for such an early increase in our own data by linear regression within selected early time windows (Table S2). As these slopes were positive, we allowed for a first phase that last for the minimum reported value of 0.5 years. The parameters of the first phase have been difficult to identify, as the initial value, *C*_0_, and the slope parameter, *α*, are correlated. We are limited by the fact that the data are cross-sectional, and the data are variable between individuals. Accepting the strong external evidence for an initial increase, we decided to also accept the slopes estimated by the *α* parameter, even if these were not significantly different from zero.

We refrained from using the exponential initial phase proposed by Wade and Ades (1994), and Payne et al. (2020), because our linear model allowed us to easily define a fixed end-point of the initial phase, and even this simple linear slope during a fixed first phase was difficult to identify. Thus, the extension of previous models with a fixed initial linear phase kept the model simple, while nevertheless allowing it to capture the different initial dynamics. We think the introduction of an initial phase is important when normalising data from very young children.

Our method is novel in the way that it stays close to experimental measurement procedures and only fits truly measured data. In most cases, these are percentages that are received via flow cytometry analysis. The conventional procedure to obtain cell count data is to multiply these percentages with the counts of a reference population (like a total lymphocyte count). While count data tend to be variable, the calculated counts are even more variable because they combine the variation in the observed counts and the observed percentages. As a sanity check, we also fitted our model (1) to conventionally calculated count data (see Figure S1 and Table S3), and indeed obtain a wider 95% confidence range of the estimated parameters (compare Table 1 with Table S3). Reassuringly, the average parameter estimates are in good agreement with each other, and as a consequence both methods provide very similar normalisation functions.

Since we are aiming for normalisation functions describing a population average, the main statistical question is what the best estimate is for the unobserved cell counts. The conventional calculated count has the advantage of using the count and percentage data from the same individual. This could be important if the counts and the percentages are dependent variables. Since the measurement of counts is difficult, and count data shows more inter-individual variability than percentages, we decided to only model what is really measured. Fortunately, both fitting procedures provide very similar normalisation functions. As techniques to count cell numbers are improving, one could in the future fit all of count data directly via model (1). Normalisation functions for the percentages would then be obtained by combining the model (1) parameters of the corresponding subpopulations using model (2), as we have done in gaining the trajectories for the percentages of the subpopulations. One could even think of fitting counts and percentages simultaneously to obtain the best population-based trajectories for counts and percentages.

In conclusion, we provide age-dependent continuous reference functions for lymphocyte data of both counts and percentages in the general Dutch population.Good agreement with a German reference [Huenecke et al., 2008] suggests that they might also serve as European references, and can be used to normalise data according to age. Our analysis does not replace the representation of lymphocyte data in contiguous age intervals, which is necessary to clinically determine the immunological health status of an individual, but rather provides an additional continuous presentation of the data. Age normalisation gains importance while quantitatively assessing changes in diseases associated with the immune system. Having a standard normalisation function at hand also simplifies the comparison of figures across different age categories by correcting for the confounding age effect. In the supplementary information, we provide a tool in the form of an R Shiny app to access the normalised absolute and relative values of all lymphocyte subsets and subpopulations according to age.

## Acknowledgement

We would like to thank Kiki Tesselaar for providing us with the data, for sharing her experimental know-how and her feedback on the manuscript, Andrew Yates and Sinead Morris for the useful discussion on the methodology and their critical reading of the manuscript, and Julia Drylewicz for statistical advice.

## Authors’ contribution

JS and RJdB conceived the study. JS performed the analysis and drafted the manuscript under supervision of RJdB. JAMB, WMB and JJMvD provided the data. JAMB and JJMvD read and edited the manuscript. All authors read and approved the manuscript.

## Supplementary Tables and Figures

**Table S1:** Data composition of the different cohorts.

| **Study** | **Dutch I** | **Dutch II** | **Dutch III** |
| --- | --- | --- | --- |
| Total | 426 | 112 | 135 |
| Children (< 18 years) | 381 | 112 | 89 |
| Adults | 45 | - | 46 |
| Age range [years] | 0-60 | 0-18 | 0-62 |
| Total lymphocyte counts | x | o |  |
| CD3+ % | x |  |  |
| CD3 counts | x |  |  |
| CD3+CD4+ % | x | o |  |
| CD3+CD4+ counts | x | o | x |
| CD27+ CD45RO- CD4+ % |  | o | x |
| CD27+ CD45RO- CD4+ counts |  | o | x |
| CD45RO+ CD4+ % |  | o | x |
| CD45RO+ CD4+ counts |  | o | x |
| CD27- CD45RO- CD4+ % |  | o | x |
| CD27- CD45RO- CD4+ counts |  | o | x |
| CD3+CD8+ % | x | o |  |
| CD3+CD8+ counts | x | o | x |
| CD27+ CD45RO- CD8+ % |  | o | x |
| CD27+ CD45RO- CD8+ counts |  | o | x |
| CD45RO+ CD8+ % |  | o | x |
| CD45RO+ CD8+ counts |  | o | x |
| CD27- CD45RO- CD8+ % |  | o | x |
| CD27- CD45RO- CD8+ counts |  | o | x |

x stands for full age range, and o for only children

**Table S2:** Linear regression analyses of subsets of the data selected according to the given time window in the early years of life. The intercept is in cells per *µ*l and the slope has a dimension of cells per *µ*l per year.

|  | Total lymphocyte count |  |  |  | CD3+ T-cell count |  |  |  | CD4+ T-cell count |  |  |  | CD8+ T-cell count |  |  |  |
| --- | --- | --- | --- | --- | --- | --- | --- | --- | --- | --- | --- | --- | --- | --- | --- | --- |
|  | Intercept | p | Slope | p | Intercept | p | Slope | p | Intercept | p | Slope | p | Intercept | p | Slope | p |
| $\leq 0.5$ years | 5729 (367) | *** | 1577 (1110) | NS. | 3470 (252) | *** | 1614 (754) | * | 2425 (191) | *** | 1142 (569) | * | 1006 (87) | *** | 390 (261) | NS. |
| $\leq 1$ year | 5984 (266) | *** | 428 (477) | NS. | 3746 (192) | *** | 469 (344) | NS. | 2670 (138) | *** | 199 (246) | NS. | 1072 (62) | *** | 100 (111) | NS. |
| $\leq 2$ years | 6228 (213) | *** | -281 (263) | NS. | 3967 (156) | *** | -162 (195) | NS. | 2863 (110) | *** | -318 (135) | * | 1084 (49) | *** | 45 (61) | NS. |
| $\leq 5$ years | 6443 (147) | *** | -655 (93) | *** | 4116 (111) | *** | -420 (76) | *** | 2895 (74) | *** | -392 (47) | *** | 1148 (35) | *** | -66 (22) | ** |
| All | 5534 (102) | *** | -138 (9) | *** | 3613 (79) | *** | -78 (7) | *** | 2356 (53) | *** | -60 (5) | *** | 1056 (23) | *** | -22 (2) | *** |

Intercept and slope estimates with standard error (SE)
F-test p-value (p) to a significant level $\geq 0.05$ , NS.; $< 0.05$ , \*; $< 0.01$ , \*\*; $< 0.001$ , \*\*\*

**Table S3:** Parameter estimates and their 95% confidence intervals obtained by fitting the calculated count data directly. Counts were calculated by multiplying the observed percentages with the observed TLC. The 95% confidence intervals for each parameter are received from bootstrapping the data 1000 times. The delay *T* was fixed to 0.5 years. The dimension of the parameters is the same as in Table 1.

| Subset | | $C_0$ | $\alpha$ | $\lambda$ | $A$ |
| --- | --- | --- | --- | --- | --- |
| CD3+ T-cell count | Estimate | 3154.9 | 1493.8 | 0.289 | 1283.6 |
|  | 95% CI | [2620.4 - 3727.2] | [115.1 - 2801.9] | [0.211 - 0.396] | [1164.5 - 1404.0] |
| CD4+ T-cell count | Estimate | 2143.1 | 978.0 | 0.384 | 858.8 |
|  | 95% CI | [1721.5 - 2527.3] | [0.0 - 2015.7] | [0.280 - 0.535] | [796.4 - 920.2] |
| CD8+ T-cell count | Estimate | 904.7 | 269.0 | 0.129 | 392.3 |
|  | 95% CI | [764.4 - 1025.4] | [0.0 - 585.9] | [0.095 - 0.173] | [338.6 - 439.0] |

**Figure S1:**
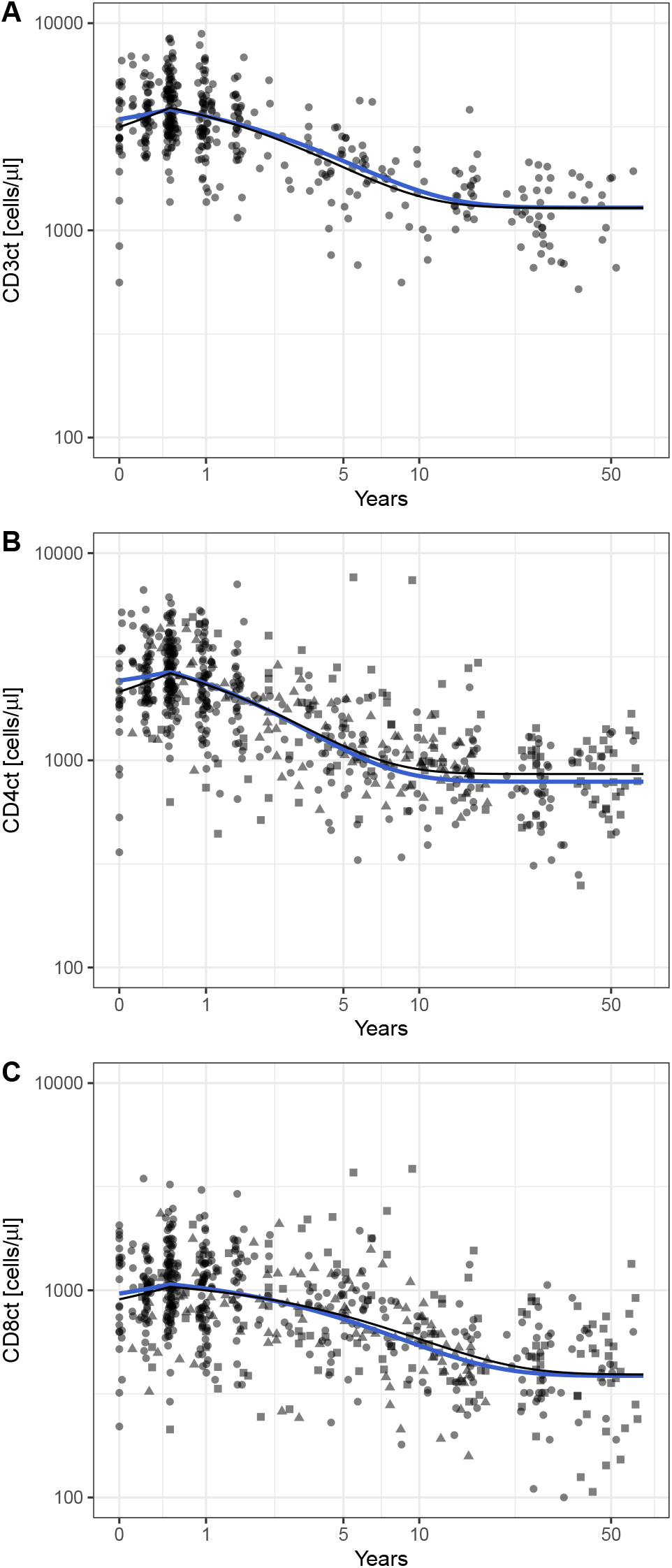
Direct fits of calculated count data are within confidence ranges of the predicted trajectories. The direct fits to the calculated CD3+ (A), CD4+ (B), CD8+ (C) cell counts are depicted by the black lines. In blue we depict our best predictions (Table 1). Bullets (•) represent data from Dutch I, triangles (▲) data from Dutch II, and squares (▪) data from Dutch III.

**Figure S2:**
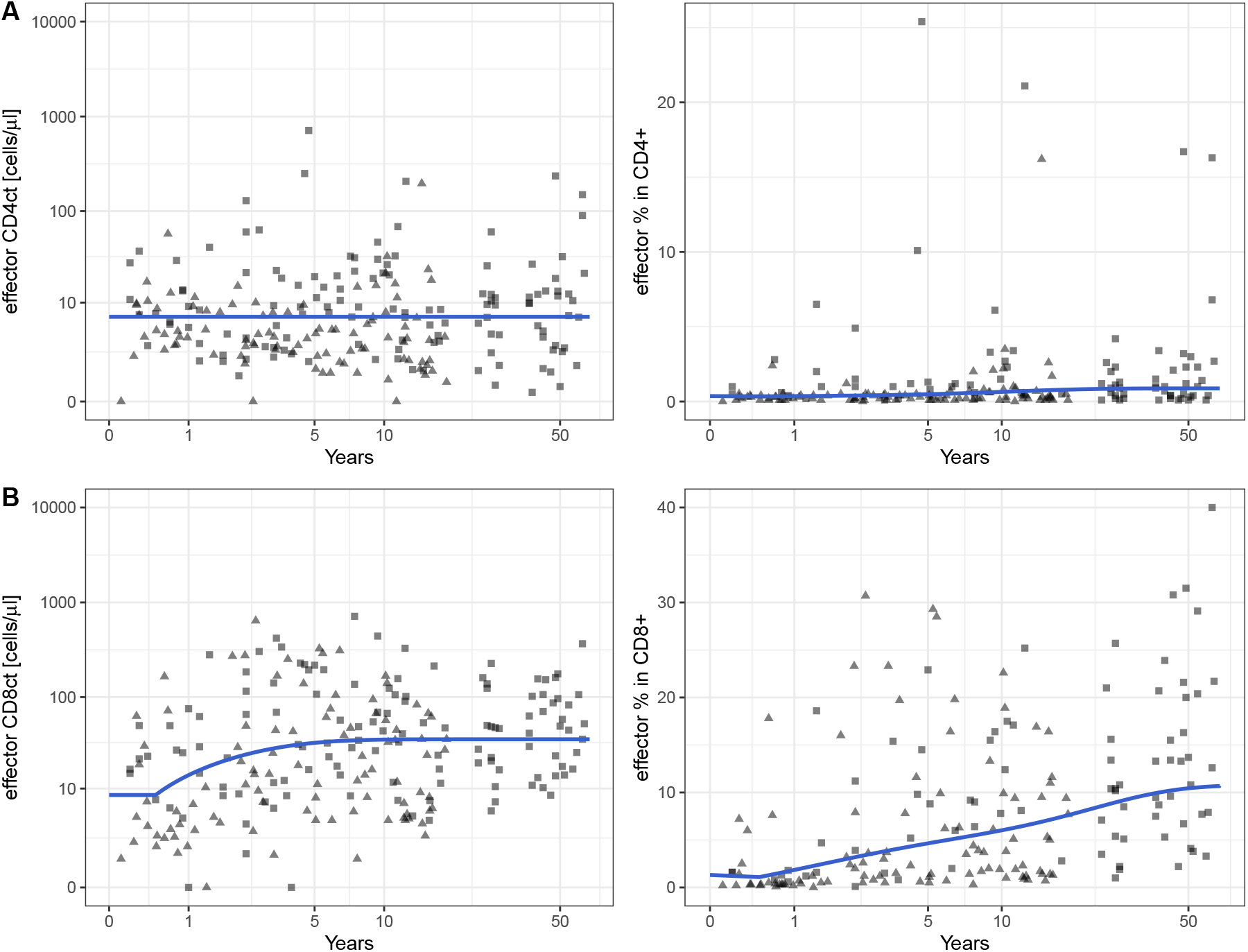
Trajectories of the effector subpopulation of CD4+ (A) and CD8+ cells (B) over age. The left panels give the absolute counts and and the right panels the percentages. Data from Dutch II are represented by triangles (▴), and data from Dutch III by squares (▪). In the left panels, the blue solid lines represent the best fits via model (1) and with the parameters presented in Table 1. The predicted trajectories for the percentages are depicted by the blue solid lines in the right panels and calculated according to model (2) where the sum of naive, memory and effector cell counts add up to 100%.

## References

K.H. Ásbjörnsdóttir, J. P. Hughes, D. Wamalwa, A. Langat, J. A. Slyker, H. M. Okinyi, J. Overbaugh, S. Benki-Nugent, K. Tapia, E. Maleche-Obimbo, A. Rowhani-Rahbar, and G. John-Stewart. Differences in virologic and immunologic response to antiretroviral therapy among HIV-1-infected infants and children. Aids, 30(18):2835–2843, nov 2016. ISSN 14735571. doi: 10.1097/QAD.0000000000001244.

I. Bains, R. Antia, R. Callard, and A. J. Yates. Quantifying the development of the peripheral naive CD4+ T-cell pool in humans. Blood, 113(22):5480–5487, 2009. doi: 10.1182/blood-2008-10.

E. Blanco, M. Pérez-Andrés, S. Arriba-Méndez, T. Contreras-Sanfeliciano, I. Criado, O. Pelak, A. Serra-Caetano, A. Romero, N. Puig, A. Remesal, J. Torres Canizales, E. López-Granados, T. Kalina, A. E. Sousa, M. van Zelm, M. van der Burg, J. J. van Dongen, and A. Orfao. Age-associated distribution of normal B-cell and plasma cell subsets in peripheral blood. Journal of Allergy and Clinical Immunology, 141(6): 2208–2219.e16, 2018. ISSN 10976825. doi: 10.1016/j.jaci.2018.02.017.

W. M. Comans-Bitter, R. De Groot, R. Van den Beemd, H. J. Neijens, W. C. Hop, K. Groeneveld, H. Hooijkaas, and J. J. Van Dongen. Immunophenotyping of blood lymphocytes in childhood: Reference values for lymphocyte subpopulations. Journal of Pediatrics, 130(3):388–393, 1997. ISSN 00223476. doi: 10.1016/S0022-3476(97)70200-2.

E. de Vries, S. de Bruin-Versteeg, M. W. Comans-Bitter, and J. J. M. van Dongen. Longitudinal follow-up of blood lymphocyte subpopulations from birth to 1 year of age. (October):586–588, 1998.

M. Goicoechea and R. Haubrich. CD4 lymphoctye percentage versus absolute CD4 lymphocyte count in predicting HIV disease progression: An old debate revisited. Journal of Infectious Diseases, 192(6):945–947, 2005. ISSN 00221899. doi: 10.1086/432972.

S. Huenecke, M. Behl, C. Fadler, S. Y. Zimmermann, K. Bochennek, L. Tramsen, R. Esser, D. Klarmann, M. Kamper, A. Sattler, D. Von Laer, T. Klingebiel, T. Lehrnbecher, and U. Koehl. Age-matched lymphocyte subpopulation reference values in childhood and adolescence: Application of exponential regression analysis. European Journal of Haematology, 80(6):532–539, 2008. ISSN 09024441. doi: 10.1111/j.1600-0609.2008.01052.x.

F. Hulstaert, I. Hannet, V. Deneys, V. Munhyeshuli, T. Reichert, M. De Bruyere, and K. Strauss. Age-related changes in human blood lymphocyte subpopulations. II. Varying kinetics of percentage and absolute count measurements, 1994. ISSN 00901229.

D. Lawrie, H. Payne, M. Nieuwoudt, and D. K. Glencross. Observed full blood count and lymphocyte subset values in a cohort of clinically healthy South African children from a semi-informal settlement in Cape Town. South African Medical Journal, 105 (7):589–595, 2015. ISSN 02569574. doi: 10.7196/SAMJnew.7914.

J. Lewis, H. Payne, A. Sarah Walker, K. Otwombe, D. M. Gibb, A. G. Babiker, R. Panchia, M. F. Cotton, A. Violari, N. Klein, and R. E. Callard. Thymic output and CD4 T-cell reconstitution in HIV-infected children on early and interrupted antiretroviral treatment: Evidence from the children with HIV early antiretroviral therapy trial. Frontiers in Immunology, 8(SEP), 2017. ISSN 16643224. doi: 10.3389/fimmu.2017.01162.

K. Li, Y. G. Peng, R. H. Yan, W. Q. Song, X. X. Peng, and X. Ni. Age-dependent changes of total and differential white blood cell counts in children. Chinese medical journal, 133(16):1900–1907, 2020. ISSN 25425641. doi: 10.1097/CM9.0000000000000854.

H. Payne, D. Lawrie, M. Nieuwoudt, M. F. Cotton, D. M. Gibb, A. Babiker, D. Glencross, and N. Klein. Comparison of Lymphocyte Subset Populations in Children From South Africa, US and Europe. Frontiers in Pediatrics, 8(July):1–12, 2020. ISSN 22962360. doi: 10.3389/fped.2020.00406.

M. Q. Picat, J. Lewis, V. Musiime, A. Prendergast, K. Nathoo, A. Kekitiinwa, P. Nahirya Ntege, D. M. Gibb, R. Thiebaut, A. S. Walker, N. Klein, R. Callard, P. Munderi, P. Nahirya-Ntege, R. Katuramu, L. Matama, F. Nankya, G. Nabulime, A. Ruberantwari, R. Sebukyu, G. Tushabe, D. Nakitto-Kesi, P. Mugyenyi, V. Musiime, R. Keishanyu, V. D. Afayo, J. Bwomezi, J. Byaruhanga, P. Erimu, C. Karungi, H. Kizito, W. S. Namala, J. Namusanje, R. Nandugwa, T. K. Najjuko, E. Natukunda, M. Ndigendawani, S. O. Nsiyona, K. Robinah, B. Bainomuhwezi, D. Sseremba, J. Tezikyabbiri, C. S. Tumusiime, A. Balaba, A. Mugumya, K. J. Nathoo, M. F. Bwakura-Dangarembizi, E. Chidziva, T. Mhute, T. Vhembo, R. Mandidewa, D. Nyoni, G. C. Tinago, J. Bhiri, D. Muchabaiwa, M. Phiri, V. Masore, C. C. Marozva, S. J. Maturure, S. Tsikirayi, L. Munetsi, K. M. Rashirai, J. Steamer, R. Nhema, W. Bikwa, B. Tambawoga, E. Mufuka, A. Kekitiinwa, P. Musoke, S. Bakeera-Kitaka, R. Namuddu, P. Kasirye, A. Babirye, J. Asello, S. Nakalanzi, N. C. Ssemambo, J. Nakafeero, J. Tikabibamu, G. Musoba, J. Ssanyu, M. Kisekka, D. M. Gibb, M. J. Thomason, A. S. Walker, A. D. Cook, B. Naidoo, M. J. Spyer, C. Male, A. J. Glabay, L. K. Kendall, and A. Prendergast. Predicting Patterns of Long-Term CD4 Reconstitution in HIV-Infected Children Starting Antiretroviral Therapy in Sub-Saharan Africa: A Cohort-Based Modelling Study. PLoS Medicine, 2013. ISSN 15491277. doi: 10.1371/journal.pmed.1001542.

R Development Core Team. R: A language and environment for statistical computing, 2003. URL http://www.r-project.org.

E. Salzmann-Manrique, M. Bremm, S. Huenecke, M. Stech, A. Orth, M. Eyrich, A. Schulz, R. Esser, T. Klingebiel, P. Bader, E. Herrmann, and U. Koehl. Joint modeling of immune reconstitution post haploidentical stem cell transplantation in pediatric patients with acute leukemia comparing CD34+-selected to CD3/CD19-depleted grafts in a retrospective multicenter study. Frontiers in Immunology, 9 (AUG):1–12, 2018. ISSN 16643224. doi: 10.3389/fimmu.2018.01841.

J. Schröter, A. J. Anelone, A. J. Yates, and R. J. de Boer. Time to Viral Suppression in Perinatally HIV-Infected Infants Depends on the Viral Load and CD4 T-Cell Percentage at the Start of Treatment. Journal of acquired immune deficiency syndromes (1999), 83(5):522–529, 2020. ISSN 19447884. doi: 10.1097/QAI.0000000000002291.

W. T. Shearer, H. M. Rosenblatt, R. S. Gelman, R. Oyomopito, S. Plaeger, E. Stiehm, D. W. Wara, S. D. Douglas, K. Luzuriaga, E. J. McFarland, R. Yogev, M. H. Rathore, W. Levy, B. L. Graham, and S. A. Spector. Lymphocyte subsets in healthy children from birth through 18 years of age. Journal of Allergy and Clinical Immunology, 112 (5):973–980, 2003. ISSN 00916749. doi: 10.1016/j.jaci.2003.07.003.

K. Soetaert and T. Petzoldt. Inverse modelling, sensitivity and monte carlo analysis in R using package FME. Journal of Statistical Software, 33(3):1–28, 2010. ISSN 15487660. doi: 10.18637/jss.v033.i03.

R. Valiathan, M. Ashman, and D. Asthana. Effects of Ageing on the Immune System: Infants to Elderly. Scandinavian Journal of Immunology, 83(4):255–266, 2016. ISSN 13653083. doi: 10.1111/sji.12413.

D. van den Heuvel, M. A. Jansen, K. Nasserinejad, W. A. Dik, E. G. van Lochem, L. E. Bakker-Jonges, H. Bouallouch-Charif, V. W. Jaddoe, H. Hooijkaas, J. J. van Dongen, H. A. Moll, and M. C. van Zelm. Effects of nongenetic factors on immune cell dynamics in early childhood: The Generation R Study. Journal of Allergy and Clinical Immunology, 139(6):1923–1934.e17, 2017. ISSN 10976825. doi: 10.1016/j.jaci.2016.10.023.

J. J. Van Dongen, M. Van Der Burg, T. Kalina, M. Perez-Andres, E. Mejstrikova, M. Vlkova, E. Lopez-Granados, M. Wentink, A. K. Kienzler, J. Philippé, A. E. Sousa, M. C. Van Zelm, E. Blanco, and A. Orfao. EuroFlow-based flowcytometric diagnostic screening and classification of primary immunodeficiencies of the lymphoid system. Frontiers in Immunology, 10(JUN):1–21, 2019. ISSN 16643224. doi: 10.3389/fimmu.2019.01271.

R. van Gent, C. M. van Tilburg, E. E. Nibbelke, S. A. Otto, J. F. Gaiser, P. L. Janssens-Korpela, E. A. Sanders, J. A. Borghans, N. M. Wulffraat, M. B. Bierings, A. C. Bloem, and K. Tesselaar. Refined characterization and reference values of the pediatric T-and B-cell compartments. Clinical Immunology, 133 (1):95–107, 2009. ISSN 15216616. doi: 10.1016/j.clim.2009.05.020. URL http://dx.doi.org/10.1016/j.clim.2009.05.020.

E. Veel, L. Westera, R. van Gent, L. Bont, S. Otto, B. Ruijsink, H. H. Rabouw, T. Mudrikova, A. Wensing, A. I. Hoepelman, J. A. Borghans, and K. Tesselaar. Impact of aging, cytomegalovirus infection, and long-term treatment for human immunodeficiency virus on CD8+ T-Cell subsets. Frontiers in Immunology, 9(MAR), 2018. ISSN 16643224. doi: 10.3389/fimmu.2018.00572.

A. M. Wade and A. E. Ades. Age-related reference ranges: Significance tests for models and confidence intervals for centiles. Statistics in Medicine, 13(22), 1994. ISSN 10970258. doi: 10.1002/sim.4780132207.

